# ShinyArchR.UiO: User-friendly, integrative and open-source tool for visualisation of single-cell ATAC-seq data using ArchR

**DOI:** 10.1101/2021.06.21.449316

**Authors:** Ankush Sharma, Akshay Akshay, Marie Rogne, Ragnhild Eskeland

## Abstract

**Motivation:** Mapping of chromatin accessibility landscapes in single-cells and the integration with gene expression enables a better understanding of gene regulatory mechanisms defining cell identities and cell-fate determination in development and disease. Generally, raw data generated from single-cell Assay for Transposase-Accessible Chromatin sequencing (scATAC-seq) are deposited in repositories that are inaccessible due to lack of in-depth knowledge of computational programming.

**Results:** We have developed ShinyArchR.UiO, an R-based shiny app, that facilitates scATAC-seq data accessibility and visualisation in a user-friendly, interactive, and open-source web interface. ShinyArchR.UiO is a tool that can streamline collaborative efforts for interpretation of massive chromatin accessible data and promotes open access data sharing for wider audiences.

**Availability and implementation:** ShinyArchR.UiO is available at https://Github.com/EskelandLab/ShinyArchRUiO and a demo server set up with a haematopoietic tutorial dataset: https://cancell.medisin.uio.no/ShinyArchR.UiO

**Contact:** &

## 1. Introduction

Transposase-accessible chromatin high throughput sequencing (ATAC-seq) is a powerful method for the assessment of genome-wide chromatin accessibility (Buenrostro *et al*., 2013) and can be explored at a single cell resolution (Cusanovich *et al*., 2015; Buenrostro *et al*., 2015). A variety of software packages exist, but no uniform analysis has been developed for scATAC-seq data (Baek and Lee, 2020). Analysis of Regulatory Chromatin in R (ArchR) is a comprehensive package for scATAC-seq analysis (Granja *et al*., 2021) that allows for integration with single-cell RNA-seq using Seurat (Stuart *et al*., 2019; Satija *et al*., 2015). Tools for sharing scATAC-seq data are currently very scarce, and there are no open-source tools based on the analysis performed using the ArchR software toolkit. ShinyArchR.UiO is a tool written in R, that adapts pre-processed ArchR data for online visualisation and sharing.

## 2. Workflow and Outputs

We have made the source code for ShinyArchR.UiO open to the public and the code can easily be customised for other scATAC-seq applications. The plots generated can be saved as high-quality pdfs.

## 3. Visualization of scATAC-seq data

We have used a down sampled tutorial dataset (Granja *et al*., 2021) to illustrate the features of ShinyArchR.UiO (Fig. 1 and S1).

**Figure 1.**
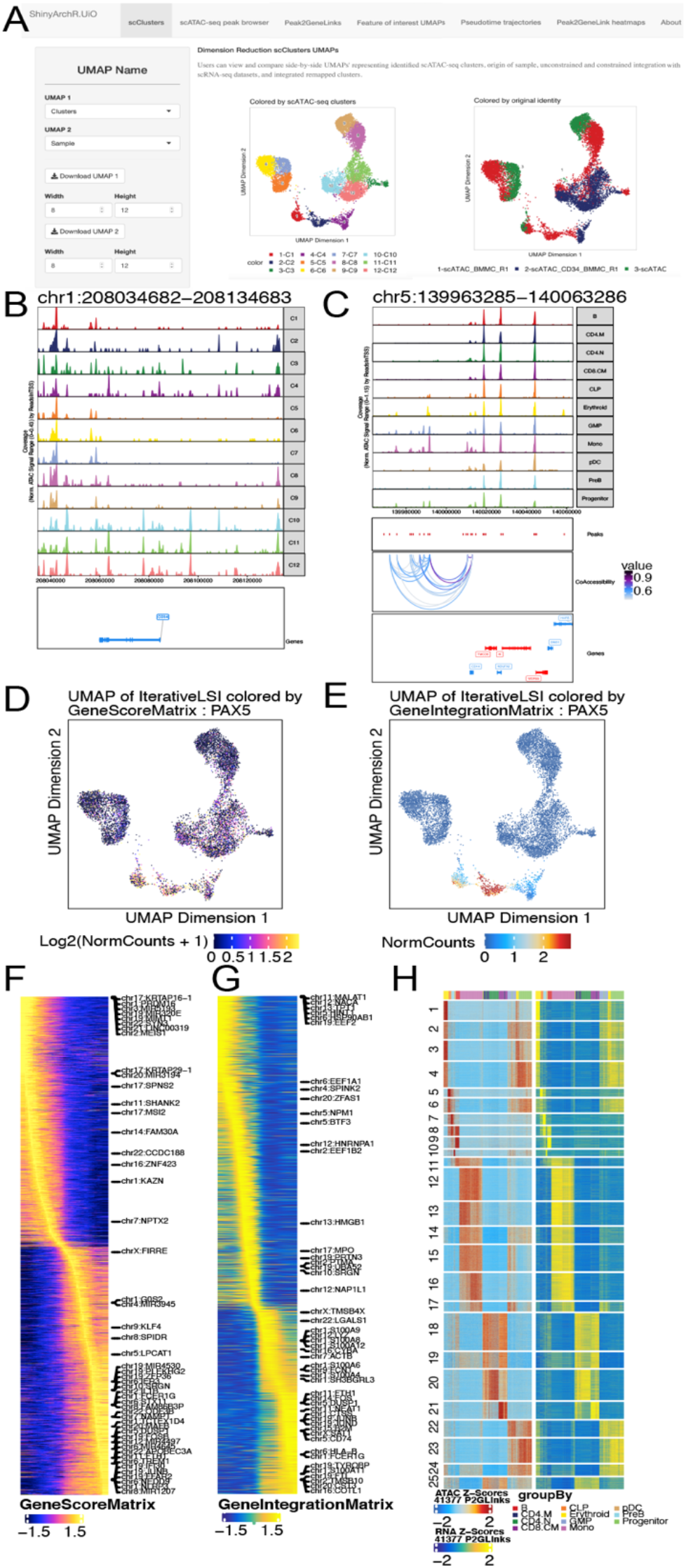
ShinyArchR.UiO have different web interfaces in tabs where the user can compute and plot scATAC-seq data from pre-processed ArchR arrow files. (a) Users can select and explore five different UMAPs side-by-side. Here shown with data from a down sampled tutorial dataset comprising scATAC-seq and scRNA-seq from hematopoietic cells. Multiple ArchR features can be plotted, viewed and saved as pdfs: (b) PlotBrowser; (c) Peak2GeneLinks browser plots; side-by-side feature comparisons of (d) GeneScoreMatrix and (e) GeneIntegrationMatrix; trajectory heatmaps including (f) GeneScoreMatrix and (g) GeneIntegrationMatrix; and (h) side-by-side heatmaps of gene scores and gene expression. The about section contains information and all references for this tool.

### 3.1 Key features of ShinyArchR.UiO

The user can:

1. Perform multidimensional reduction UMAP plot of original samples, scATAC-seq clusters, and clusters from cross-platform linkage integration with scRNA-seq generated by ArchR; 2. Explore plot browser peaks grouped by original samples or scATAC-seq clusters with an adjustable window of 250 KB distance from the centre; 3. View co-accessibility of peaks from clusters based on Peak2GeneLink analysis; 4. Compare GeneScore and Geneintegration matrix UMAPs for selected genes side-by-side; 5. Visualize four different ArchR defined pseudotime trajectory heatmaps; and 6. Explore Peak2GeneLink heatmap on scATAC-seq and scRNA-seq modality.

### 3.2 Basic usage

ShinyArchR.UiO uses pre-processed ArchR HDF5 formatted objects stored on disk and generates the ShinyArchR.UiO for visualization of the scATAC-seq metadata. Retrieved information is plotted in ShinyArchR.UiO as peaks, UMAPs and heatmaps as shown (Fig. 1). This process can be executed in two simple steps:

i. Download or git clone: https://github.com/EskelandLab/ShinyArchRUiO
ii. Provide a path to saved arrow file folders (Saved-ArchRproject) obtained from ArchR analysis in *glabal*.*R* and name given to trajectory analysis in the getTrajectory function of ArchR, see Si (2) for more details.

~~~
Running ShinyArchR.UiO from command line
R -e “shiny::run-App(‘∼/ShinyArchR.UiO’,launch.browser = TRUE)”
Or open app.R or global.R, press Run App button on R Graphical User Interface
~~~

The initiation of the shiny app takes 5-10 minutes on quad-core 16 gigabytes of RAM for data representing 10000 cells.

### 3.3 Advanced usage

ShinyArchR.UiO can be run locally or be made available to be hosted on an open-source shiny-server, offering features such as apps behind firewalls (RStudio Team). Genome-wide maps of open chromatin regions of multiple pre-processed scATAC-seq datasets can result in a large amount of data, therefore, we highly recommend evaluating the storage capacity for hosting purposes. We have made an open-source R-code tool for plotting and visualization of scATAC-seq data including integration with complementary scRNA-seq for a broader range of users than experienced computer scientists. The ShinyArchR.UiO web interface allows the user to easily explore chromatin opening across the genome at a single-cell level for improved insight into complex biological data.

## Author Contributions and Funding

AS and AA developed the code with scientific input and tests performed by MR and RE. RE and AS wrote the manuscript with input from MR. This work was partly supported by the Research Council of Norway through its Centres of Excellence funding scheme, project number 262652. Conflict of interest, none declared.

## Acknowledgments

We would like to acknowledge the Department for Research Computing, UiO for help to set up the Shiny server for demo ShinyArchR.UiO. Emily L. B. Martiensen, Hallvard A. Wæhler and Sakshi Singh for testing ShinyArchR.UiO setup on different operating systems. We thank Leslie Foster for testing ShinyArchRUiO and proofreading the manuscript.

## Supplementary information

is linked to the online version of the paper.

