## Supplemental File for "ShinyArchR.UiO: User-friendly, integrative and open-source tool for visualisation of single-cell ATAC-seq data using ArchR"

---

### Supplementary Information

## SI (1)

The Analysis of Regulatory Chromatin in R (ArchR) package is used for analysis of single-cell chromatin accessibility data and requires high-performance computing environments (Granja *et al.*, 2021). ArchR software toolkit can be utilized for scATAC-seq data obtained from multiple scATAC-seq implementations, including the 10x Genomics Chromium system. The ArchR software stores all data (i.e., metadata, data matrices, and accessible fragments) associated with each sample in Arrow files. ShinyArchR.UiO requires HDF file formatted Arrow files stored on a disk or server, which greatly reduces the memory footprint for obtaining different plots from massive-scale single-cell chromatin accessibility data.

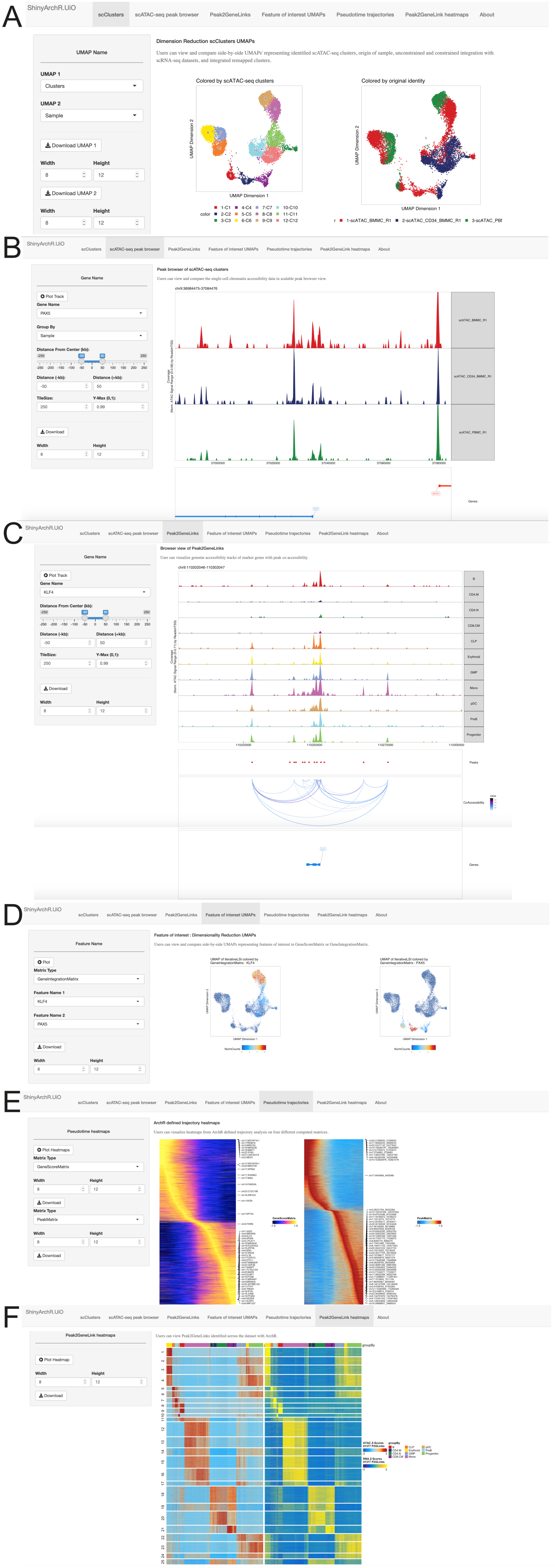


**Figure S1**. The ShinyArchR.UiO application presents six tabs where users can visualize single-cell chromatin accessibility data. A) scClusters shows multidimensional reductions UMAP plot (McInnes et al., 2020) of the single-cell data, where cell information about original identity, scATAC-seq clusters and subpopulations after integrations with scRNA-seq can be selected in the dropdown menu to be viewed side-by-side. B) The scATAC-seq peak browser tab shows chromatin accessibility data in a peak browser view. The user can plot select for individual samples or scATAC-seq clusters for genes of interest in a scalable window size and download pdfs. C) Genome accessibility browser tracks can also be plotted and viewed for selected genes with Peak2GeneLinks. D) The Feature of Interest UMAPs tab allows users to compare one or two selected genes side-by-sideon GeneScoreMatrix or GeneIntegratioMatrix. E) The Pseudotime trajectory tab allows users to visualise pseudotime trajectories heatmaps for four different matrices: GeneScore matrix, GeneIntegration matrix, Motif matrix, and Peak matrix. F) The Peak2GeneLinks heatmaps visualizes the Peak2GeneLinks identified across the scATAC-seq and scRNA-seq datasets (Gu et al., 2016). Users will find information about the scope, approach, credits, and link to Github repository in the about tab.

## SI (2)

##### A detailed tutorial for setup:

Single-cell chromatin accessibility data can be processed and analysed using ArchR (Granja *et al.*, 2021). The analysis can be performed using standard ArchR settings on a test dataset of hematopoietic cells or can be applied on user's own datasets. The project folders saved can be used as input for visualization of data by ShinyArchR.UiO: 1) Project-folder1: Cross-platform linkage of scATAC-seq cells with scRNA-seq cells analysis of ArchR analysis., 2) Project-folder2: Labelling scATAC-seq clusters with scRNA-seq information and 3) Project-folder3: Pseudotime trajectory analysis. Genome-wide maps of open chromatin regions of multiple pre-processed scATAC-seq datasets can result in a large amount of data, therefore, we highly recommend evaluating the storage capacity for hosting purposes

## SI (3)

##### Please follow the steps for ShinyArchR.UiO setup on the local system:

1. Download ShinyArchR.UiO or git clone at https://Github.com/EskelandLab/ShinyArchRUiO.
2. Provide the path to the saved folders in global.R
```
savedArchRProject1 <- loadArchRProject("path to project-folder1/")
savedArchRProject2 <- loadArchRProject("path to project-folder2/")
savedArchRProject3 <- loadArchRProject("path to project-folder3/")
```
3. Provide trajectory name as given in your analysis in getTrajectory function in chunk "Add Metadata of Trajectory" in global.R
4. Open app.R in RStudio or on R graphical User Interface, press RunApp to run ShinyArchR.UiO.
5. Setting ShinyArchR.UiO from the terminal.
   Run the command
```
R -e "shiny::runApp('~/ShinyArchR.UiO',launch.browser = TRUE)"
```

The initiation of the shiny app takes 5-10 minutes on quad-core 16 gigabytes of RAM for data representing 10000 cells.

## SI (4)

##### Note:

1. R version 4.0.0 and over is recommended.
2. Please ensure ArchR (Granja *et al.*, 2021), Seurat (Stuart *et al.*, 2019; Satija *et al.*, 2015), Magic (Dijk *et al.*, 2018), ggplot2 (Wickham, 2009), rtracklayer (Lawrence *et al.*, 2009) and other dependencies required are installed and loaded properly in the R environment. More details on the procedure to install packages and their dependencies can be found on Github readme.
   Users can raise an Github issue on the ShinyArchR.UiO GitHub repository if they face errors related to set up.

## SI (5)

##### Frequently Asked Questions

- Q: How much memory/storage space does ShinyArchR.UiO and the shiny app consume?
    a. The shiny app itself is not memory intensive and is meant to be a heavy-duty app where multiple users can access the app at the same time. The memory required is dependent on the size of saved project files from ArchR.
    b. Simultaneously, ArchR employs Arrow files, an HDF5 file format, to store massive single-cell chromatin accessibility data on disk.
    c. Initial setup of ShinyArchr.UiO is computation intensive. This includes steps for computing marker genes for Peak2Genelinks analysis and other plots. A typical laptop with 4GB RAM can handle datasets from estimated 10-20k cells while 16GB RAM machines can handle around 30k-100k cells.

- Q: Does ShinyArchR.UiO support performance gain across different operating systems?
    a. ShinyArchR.UiO is a tool to visualize scATAC seq data analysed by the ArchR software, which is predominantly optimized for Unix-based systems such as Linux and MacOS. ShinyArchR.UiO is tested and should work on Windows. However, users cannot take advantage of parallelization on Windows.
